# Cyp7b1-inhibiting azoles as novel enhancers of hematopoietic stem and progenitor cell mobilization

**DOI:** 10.1101/2025.10.13.682145

**Authors:** Brandon L. Vu, Travis J. Roeder, Jitendra K. Kanaujiya, Amy L. Kimble, Eddy Tsang, Hideyuki Oguro

**Affiliations:** Department of Cell Biology, University of Connecticut School of Medicine, Farmington, CT, USA

## Abstract

Mobilized hematopoietic stem and progenitor cells (HSPCs) are essential for transplantation-based therapies, including curative gene therapies for sickle cell disease (SCD). While granulocyte colony-stimulating factor (G-CSF, filgrastim) remains the standard mobilization agent, many patients respond inadequately, and it can trigger life-threatening vaso-occlusive crises in SCD. The CXCR4 antagonist AMD3100 (plerixafor) is routinely combined with G-CSF for non-SCD settings but is ineffective as a single agent in SCD, underscoring the urgent need for alternative strategies. We previously identified 27-hydroxycholesterol (27HC) as a physiological inducer of HSPC mobilization during pregnancy. Here, we show that exogenous 27HC enhances AMD3100-induced HSPC mobilization in mice, either alone or with G-CSF. Because 27HC is metabolized by the enzyme Cyp7b1, we tested whether pharmacological Cyp7b1 inhibition could mimic this effect. Treatment with clotrimazole, an antifungal and Cyp7b1 inhibitor, significantly enhanced AMD3100-induced HSPC mobilization in wild-type, SCD, and humanized mice. Importantly, intravenous administration of voriconazole, a clinically approved systemic antifungal with Cyp7b1-binding activity, similarly augmented AMD3100-induced HSPC mobilization in wild-type and SCD mice without altering steady-state hematopoiesis. These findings establish Cyp7b1-inhibiting azoles as novel and clinically relevant enhancers of HSPC mobilization, particularly for SCD patients who cannot safely receive G-CSF but require robust HSPC yields for gene therapy.

**Graphical abstract:** 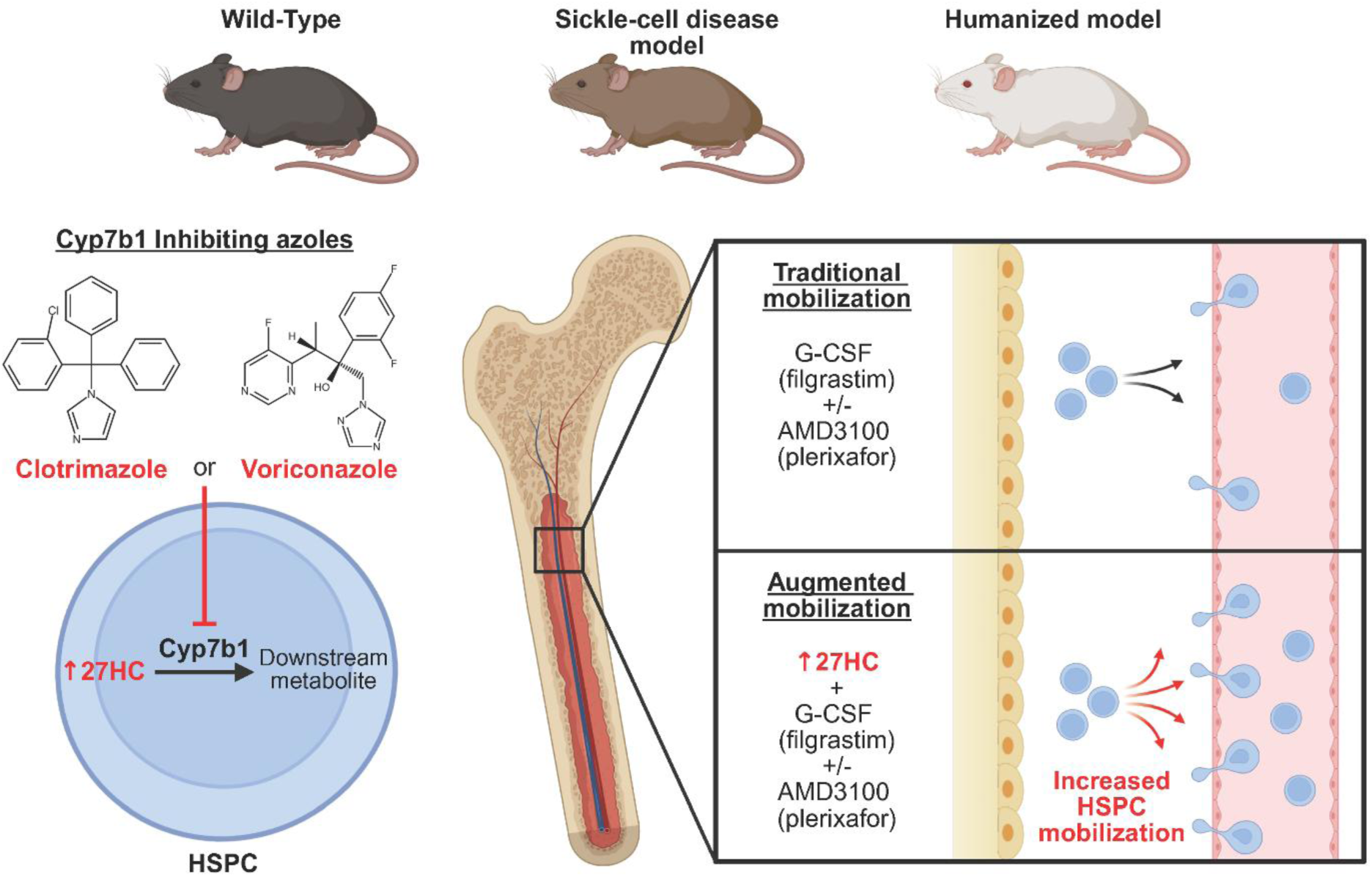

## Introduction

Adult hematopoietic stem and progenitor cells (HSPCs) reside in the bone marrow and sustain blood production throughout life (1), and HSPC collection and transplantation are central to therapies for hematologic malignancies and blood disorders (2). The most common HSPC collection strategy uses granulocyte colony-stimulating factor (G-CSF, filgrastim), which mobilizes HSPCs from the bone marrow into the peripheral blood for collection via apheresis. In donors and patients with poor responses to G-CSF alone, the CXCR4 antagonist AMD3100 (plerixafor) is added to enhance mobilization (3). Nonetheless, many donors and patients fail to mobilize adequate HSPCs with traditional mobilization regimens, leading to multiple apheresis sessions or transplant failure (4).

Sickle-cell disease (SCD) is an autosomal recessive blood disorder caused by mutations in the β-globin gene (HBB) that produces deoxygenated hemoglobin S (HbS), resulting in sickle-shaped red blood cells (5). SCD causes anemia and various types of crises, potentially leading to severe complications and premature death. Allogenic transplantation of HSPCs obtained from a healthy donor is curative (6), but this approach presents significant challenges, including the difficulty of identifying a suitable human leukocyte antigen (HLA)-matched donor, the risk of graft rejection, and the potential development of graft-versus-host disease (GVHD). Autologous transplantation with genetically modified HSPCs circumvents these barriers, and several gene therapy strategies—including lentiviral gene delivery and CRISPR genome-editing approaches— are now FDA-approved as novel therapies to cure SCD (7, 8). While these approaches show promise, successful HSPC gene therapies require the collection of large numbers of HSPCs from patients. Since G-CSF can trigger fatal vaso-occlusive crises in SCD patients (9–11), AMD3100 is used as a sole mobilizer to collect HSPCs from SCD patients. However, responses to AMD3100 alone are variable and less effective than G-CSF (12), highlighting the need for alternative HSPC mobilizing strategies.

To better understand how HSPC mobilization is regulated, we previously used pregnancy in mice as a model system and found that HSPC mobilization is naturally induced during pregnancy to replenish red blood cells through extramedullary hematopoiesis (13). These responses are dependent on the levels of 27-hydroxycholesterol (27HC), a cholesterol-derived oxysterol and endogenous estrogen receptor (ER) ligand (14, 15). We demonstrated that the treatment of mice with 27HC induces ERα-dependent HSPC mobilization and that mice deficient for Cyp27a1, a sterol hydroxylase necessary to generate 27HC from cholesterol (16), display impaired HSPC mobilization and splenic erythropoiesis during pregnancy (17).

Healthy people and autologous transplantation patients with higher low-density lipoprotein (LDL) or total cholesterol levels mobilize a larger number of HSPCs naturally or in response to G-CSF treatment, as compared with those with lower cholesterol levels (18, 19). High-density lipoprotein (HDL) cholesterol levels are negatively correlated with mobilization of HSPCs and monocytes (20, 21). Similarly, in mice, increased cholesterol levels are known to promote HSPC mobilization (22–25). Given that 27HC levels increase as cholesterol levels increase (26), these findings suggest that elevated 27HC levels promote HSPC mobilization in mice and humans.

Here, we demonstrate that exogenous 27HC supplementation enhances HSPC mobilization in combination with AMD3100 and G-CSF. Given that 27HC is the most abundant oxysterol in human plasma, we used pharmacological approaches to mimic 27HC supplementation. The cytochromes P450 enzyme Cyp7b1 catalyzes the metabolism of 27HC into a downstream metabolite via its oxysterol 7α-hydroxylase activity, and *Cyp7b1*-deficient mice exhibit elevated 27HC levels in tissues and plasma (27). We show that clotrimazole, a Cyp7b1-inhibiting antifungal (28), augments AMD3100-induced mobilization in wild-type, SCD, and humanized mice. Importantly, voriconazole, an FDA-approved systemic antifungal that also binds Cyp7b1 (29), produces similar effects without altering hematopoiesis. Our findings identify modulation of 27HC metabolism through Cyp7b1 inhibition as a novel, clinically relevant strategy to improve HSPC mobilization, particularly for SCD patients who cannot receive G-CSF but require robust HSPC yields for gene therapy.

## Results

### Exogenous 27HC supplementation enhances HSPC mobilization

We previously showed that exogenous 27HC supplementation enhances G-CSF-induced HSPC mobilization (17). To test whether 27HC supplementation also augments AMD3100-induced HSPC mobilization, wild-type C57BL/6J mice were treated with 27HC (10 mg/kg/day, subcutaneous) for two days, followed by AMD3100 (5 mg/kg, subcutaneous) one hour before blood collection. Mobilization of functional HSPCs was quantified by colony-forming assays using peripheral blood mononuclear cells. Two measures were used: colony-forming unit-cell (CFU-C), which represents HSPCs capable of generating any hematopoietic colony, and CFU-granulocyte, monocyte, erythroblast and megakaryocyte (CFU-gmEM), which represents the most primitive HSPC subset retaining multilineage differentiation potential to form colonies containing these four lineages. Co-administration of 27HC and AMD3100 significantly increased both CFU-C (2.7-fold) and CFU-gmEM (2.6-fold) in the peripheral blood compared with AMD3100 alone (**Figures 1, A and B**).

**Figure 1.**
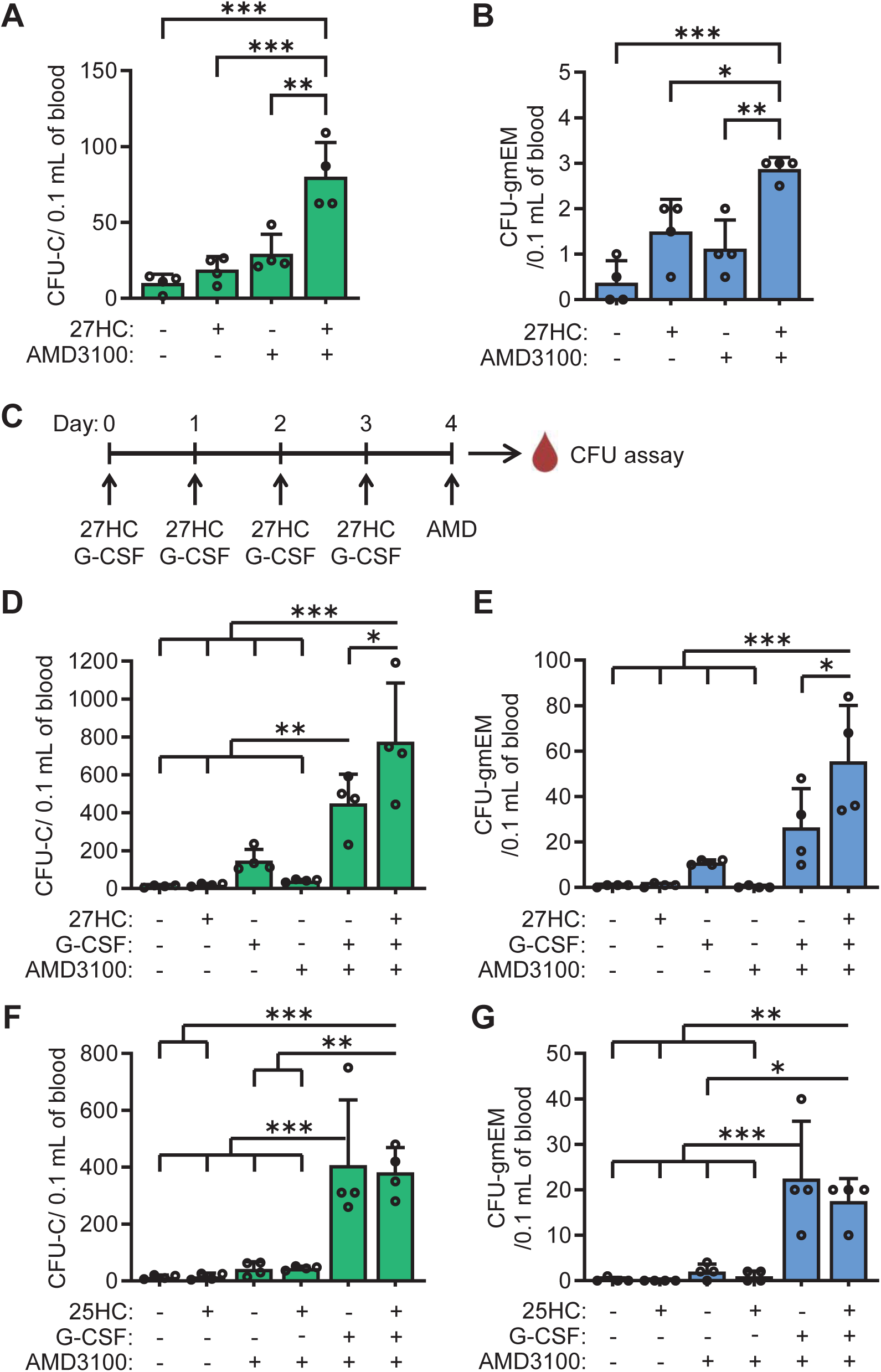
Enhanced HSPC mobilization in wild-type mice co-treated with 27HC and clinically used HSPC mobilizing agents. (**A-B**) Wild-type C57BL/6J mice were treated with 27HC or vehicle daily for two days, followed by AMD3100 (plerixafor) or vehicle one hour before peripheral blood collection for CFU assays. Numbers of CFU-C (**A**) and CFU-gmEM (**B**) after the indicated treatments (n=4 mice per condition from 3 experiments). (**C**) Experimental design for HSPC mobilization by co-treatment with 27HC, G-CSF, and AMD3100. (**D**-**E**) C57BL/6J mice were co-treated with 27HC, G-CSF and/or vehicle daily for four days, followed by AMD3100 or vehicle one hour before peripheral blood collection for CFU assays. Numbers of CFU-C (**D**) and CFU-gmEM (**E**) in peripheral blood after the indicated treatments (n=4 mice per condition from 4 experiments). (**F**-**G**) C57BL/6J mice were co-treated with 25HC, G-CSF and/or vehicle daily for four days, followed by AMD3100 or vehicle one hour before peripheral blood collection for CFU assays. Numbers of CFU-C (**F**) and CFU-gmEM (**G**) in peripheral blood after the indicated treatments (n=4 mice per condition from 3 experiments). Data represent mean ± standard deviation (SD). Statistical significance was determined by one-way ANOVA with Tukey’s multiple comparisons test (^*^P<0.05, ^**^P<0.01, ^***^P<0.001).

Because AMD3100 is used clinically when patients and donors fail to respond to G-CSF alone, we next tested whether 27HC supplementation further enhances HSPC mobilization in combination with G-CSF and AMD3100. C57BL/6J mice were treated with 27HC and/or G-CSF (250 µg/kg/day, subcutaneous) for four days, followed by AMD3100 one hour before blood collection (**Figure 1C**). Supplementation of 27HC significantly increased the numbers of mobilized CFU-C (1.7-fold) and CFU-gmEM (2.1-fold) compared with the combination of G-CSF and AMD3100 alone (**Figures 1, D and E**).

To test whether the mobilization-enhancing effect is specific to 27HC or shared with other oxysterols, we treated C57BL/6J mice with 25-hydroxycholesterol (25HC) at the same dose as 27HC. Both 27HC and 25HC can signal through estrogen receptors and liver X receptors (LXRs) (15). However, unlike 27HC, 25HC did not increase CFU-C or CFU-gmEM under either the AMD3100-only or G-CSF plus AMD3100 regimens (**Figures 1, F and G**). These findings demonstrate that elevated 27HC levels, but not 25HC levels, enhance HSPC mobilization, suggesting a 27HC specific mechanism in regulating mobilization.

### Clotrimazole enhances HSPC mobilization induced by G-CSF and AMD3100

Given that 27HC is one of the most abundant oxysterols in human plasma (30), increasing its levels by direct supplementation may be clinically challenging. The enzyme Cyp7b1 metabolizes 27HC into downstream products through its oxysterol 7α-hydroxylase activity, and *Cyp7b1*-deficient mice exhibit elevated 27HC levels in plasma and tissues (27). Notably, *Cyp7b1* is expressed at higher levels in long-term hematopoietic stem cells (LT-HSCs) compared with hematopoietic progenitors or differentiated cells (**Figures 2, A and B**) (31, 32), suggesting that LT-HSCs preferentially metabolize 27HC and thereby limit its intracellular availability.

**Figure 2.**
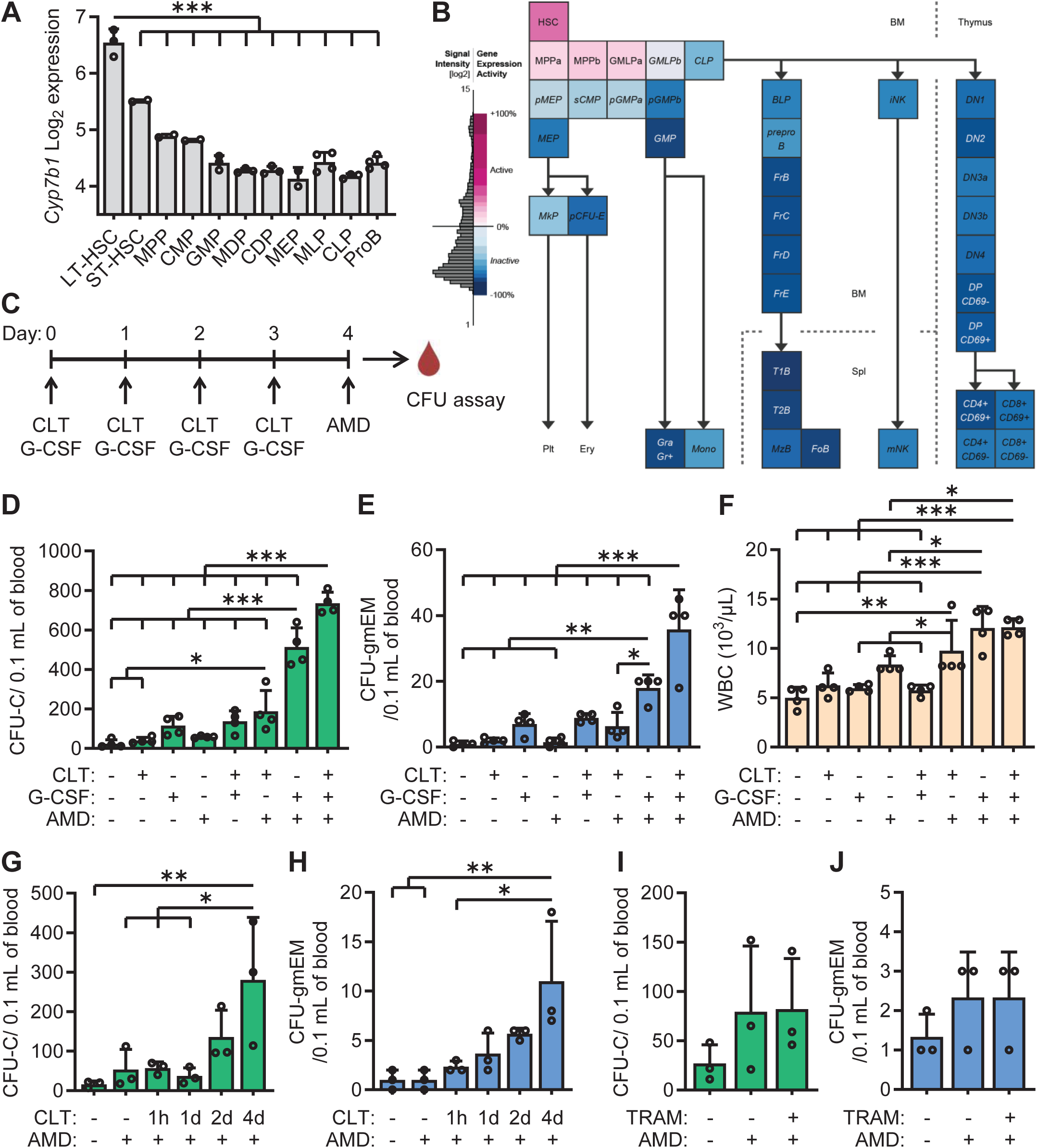
A Cyp7b1 inhibitor, clotrimazole, enhances HSPC mobilization in combination with clinically used mobilizing agents. (A) Expression of *Cyp7b1* mRNA in bone marrow hematopoietic cells. Data from BloodSpot (31), mouse immugen stem and progenitor cell dataset (n=2-4) (51). LT34F, ST34F, MPP34F, proB.CLP, and proB.FrA were converted to LT-HSC, ST-HSC, MPP, CLP, and ProB, respectively. (B) Expression of *Cyp7b1* mRNA in hematopoietic cells from bone marrow, spleen, and thymus, generated using Gene Expression Commons (mouse hematopoiesis model, probeset 1421074_at) (32). (C) Experimental design for HSPC mobilization by co-treatment with clotrimazole (CLT), G-CSF, and AMD3100. (**D**-**F**) C57BL/6J mice were co-treated with CLT, G-CSF and/or vehicle daily for four days, followed by AMD3100 or vehicle one hour before peripheral blood collection for CFU assays. Numbers of CFU-C (**D**) and CFU-gmEM (**E**) in peripheral blood and white blood cell (WBC) counts (**F**) after the indicated treatments (n=4 mice per condition from 4 experiments). (**G**-**H**) C57BL/6J mice were treated with CLT or vehicle daily for the indicated times, followed by AMD3100 or vehicle one hour before peripheral blood collection for CFU assays. Numbers of CFU-C (**G**) and CFU-gmEM (**H**) in the peripheral blood (n=3 mice per condition from 3 experiments). (**I**-**J**) C57BL/6J mice were treated with TRAM-34 (TRAM), an IK_Ca_ channel blocker, or vehicle daily for four days, followed by AMD3100 or vehicle one hour before peripheral blood collection for CFU assays. Numbers of CFU-C (**I**) and CFU-gmEM (**J**) in the peripheral blood (n=3 mice per condition from 3 experiments). Data represent mean ± SD. Statistical significance was determined by one-way ANOVA with Tukey’s multiple comparisons test (^*^P<0.05, ^**^P<0.01, ^***^P<0.001).

We used clotrimazole, an antifungal agent known to inhibit Cyp7b1 function (28), to test whether pharmacologic inhibition of Cyp7b1 elevates intracellular 27HC levels in HSPCs and enhances their mobilization. In a similar mobilization regimen (**Figure 2C**), C57BL/6J mice co-treated with clotrimazole (50 mg/kg/day, intraperitoneally for four days) and AMD3100 displayed greater numbers of mobilized CFU-C (3.2-fold) and CFU-gmEM (4.3-fold) compared with AMD3100 alone (P=0.02 and P=0.04, respectively; one-way ANOVA across vehicle, clotrimazole, AMD3100, and co-treatment of clotrimazole and AMD3100 groups) (**Figures 2, D and E**). While clotrimazole combined with G-CSF did not significantly increase mobilized CFU-C or CFU-gmEM compared with G-CSF alone, clotrimazole significantly increased mobilized CFU-C (1.4-fold) and CFU-gmEM (2.0-fold) when co-administered with G-CSF and AMD3100 (**Figures 2, D and E**).

Although the combination of G-CSF and AMD3100 significantly increased white blood cell (WBC) counts compared with vehicle control, the addition of clotrimazole did not further increase WBCs (**Figure 2F**). This suggests that clotrimazole preferentially enhances mobilization of immature HSPCs rather than mature leukocytes, consistent with the restricted expression of its target, Cyp7b1, in LT-HSCs.

We next tested whether shorter dosing regimens were sufficient to enhance HSPC mobilization. C57BL/6J mice were treated with clotrimazole for one hour, one day, two days, or four days, followed by AMD3100 for one hour. While two days of clotrimazole treatment showed a trend toward enhanced mobilization of CFU-C (2.5-fold) and CFU-gmEM (5.7-fold), only the four-day regimen produced a significant increase in CFU-C (5.3-fold) and CFU-gmEM (11-fold) compared with AMD3100 alone (**Figures 2, G and H**).

Clotrimazole is also known to inhibit the intermediate-conductance Ca^2+^-activated K^+^ (IK_Ca_ or Gardos) channel, thereby reducing erythrocyte dehydration in SCD mice and patients (33, 34). To determine whether IK_Ca_ channel inhibition contributed to enhanced HSPC mobilization, we treated mice with TRAM-34, a clotrimazole analog that selectively blocks IK_Ca_ channel without inhibiting cytochrome P450 activity (35). Unlike clotrimazole, IK_Ca_ channel inhibition by TRAM-34 did not promote AMD3100-induced HSPC mobilization (**Figures 2, I and J**). Together, these findings indicate that clotrimazole treatment mimics the effects of 27HC supplementation by enhancing HSPC mobilization in conjunction with clinically used HSPC mobilizers, primarily through inhibition of Cyp7b1 rather than blockade of the IK_Ca_ channel.

### Clotrimazole enhances human HSPC mobilization in humanized mice

To examine the effect of clotrimazole on human HSPC mobilization, we used a humanized mouse model transplanted with cord blood-derived human HSPCs. CD34^+^ HSPCs were magnetically enriched from freshly obtained cord blood from three independent donors, and cells from each donor were transplanted into unconditioned NBSGW immunodeficient mice (36). Four months post-transplantation, humanized mice were treated with clotrimazole and/or AMD3100, followed by peripheral blood colony assays and flow cytometric analysis of the bone marrow (**Figure 3A**). For the colony assays, methylcellulose medium supplemented exclusively with human cytokines was used, ensuring selective support of colonies derived from human HSPCs. Generated colonies expressed either human CD33 (hCD33, granulocyte/monocyte marker), hCD235a (erythroid marker), or both (**Figure 3B**). Co-treatment with clotrimazole and AMD3100 significantly increased the number of mobilized human CFU-C in peripheral blood compared with vehicle-treated controls (4.4-fold) and showed a strong trend toward increased mobilization compared with AMD3100 single treatment (2.8-fold) (**Figure 3C**). Because human HSPC chimerism varied across recipients, we normalized the levels of humanization by calculating the peripheral blood CFU-C number per bone marrow human HSPC, defined as mouse CD45 (mCD45)^-^hCD45^+^HLA-ABC^+^hCD34^+^hCD38^-^ (**Figure 3D**). After adjustment, clotrimazole co-administration significantly enhanced AMD3100-induced mobilization of human HSPCs by 2.4-fold (**Figure 3E**). These results demonstrate that pharmacologic inhibition of Cyp7b1 can augment human HSPC mobilization in vivo, highlighting its potential as a translational strategy to improve HSPC collection for clinical transplantation.

**Figure 3.**
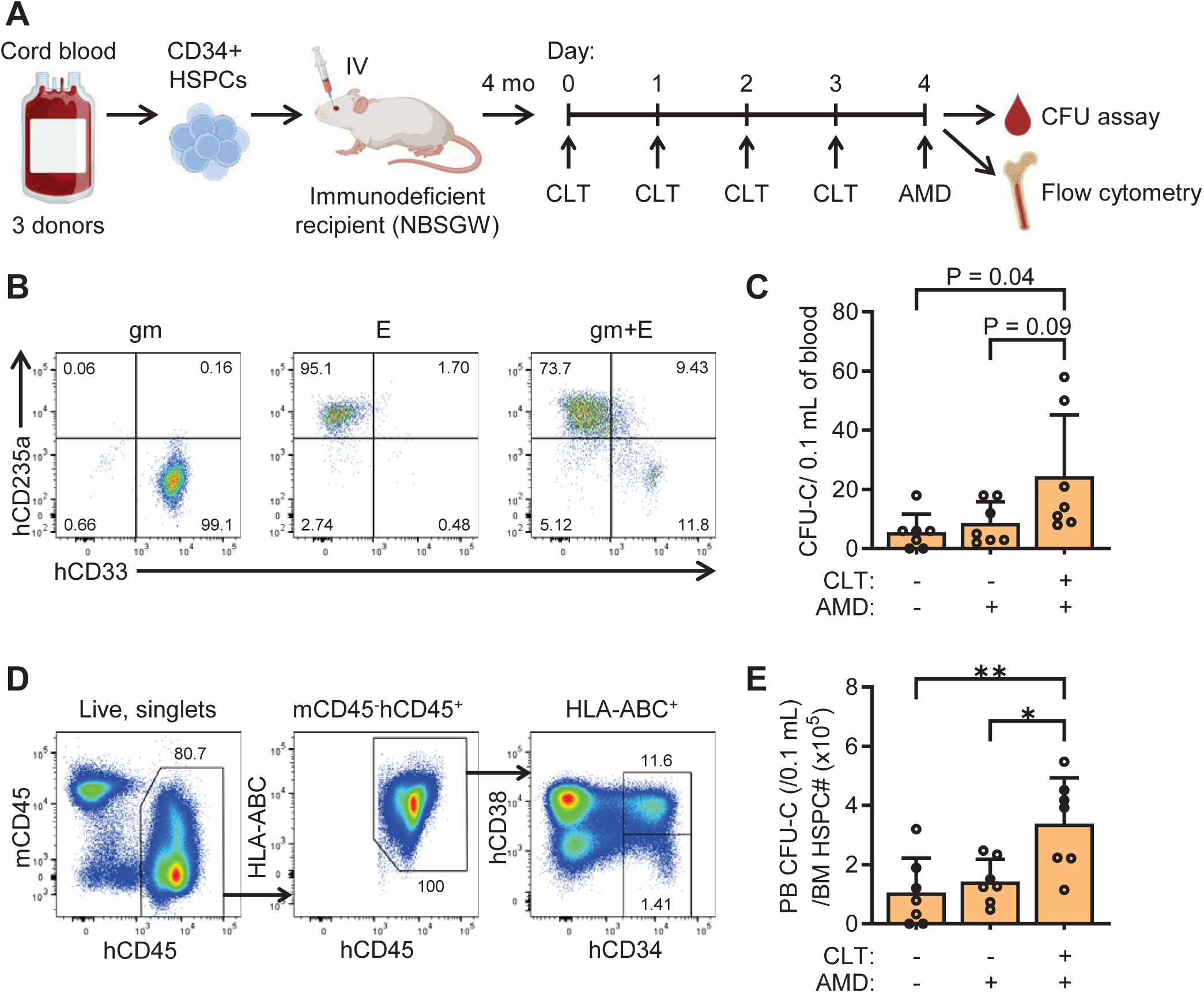
Clotrimazole enhances human HSPC mobilization in immune-humanized mice co-treated with AMD3100. (A) Immunodeficient NBSGW mice were transplanted with CD34^+^ HSPCs from human cord blood. After four months of engraftment, humanized mice were treated with CLT or vehicle daily for four days, followed by AMD3100 or vehicle one hour before analysis. (B) Flow cytometry dot plots of representative colonies derived from peripheral blood of humanized mice. Human granulocytes/macrophages (gm, hCD33^+^hCD235a^-^), erythroblasts (E, hCD235a^+^hCD33^-^), or mixed colonies (gm+E) are shown. (C) Absolute numbers of mobilized human CFU-C in peripheral blood of humanized mice after the indicated treatments (n=7 mice per condition from 4 experiments). (D) Representative flow cytometry dot plots of mCD45^-^hCD45^+^HLA-ABC^+^CD34^+^CD38^-^ human HSPCs in bone marrow of humanized mice. (E) Ratio of mobilized human CFU-C in peripheral blood (PB) to the number of bone marrow (BM) human HSPCs in humanized mouse (n=7 mice per condition from 4 experiments). Data represent mean ± SD. Statistical significance was determined by one-way ANOVA with Tukey’s multiple comparisons test (^*^P<0.05, ^**^P<0.01).

### Clotrimazole enhances HSPC mobilization in sickle cell disease mice

We next examined whether clotrimazole could enhance HSPC mobilization in the “Townes” SCD mouse model (37). Because G-CSF is contraindicated for HSPC mobilization in SCD patients due to the risk of fatal vaso-occlusive crises (9–11), we focused on testing the potential synergy between clotrimazole and AMD3100 (**Figure 4A**). Vehicle-treated SCD mice homozygous for human HbS knock-in alleles (S/S) exhibited markedly elevated baseline levels of mobilized CFU-C (15-fold, P=0.01, unpaired t test with Welch’s correction) and CFU-gmEM (16-fold, P=0.02) in the peripheral blood compared with vehicle-treated wild-type C57BL/6J mice (**Figures 4, B and C**). Although AMD3100 alone had little effect on HSPC mobilization in wild-type C57BL/6J mice (**Figures 1 and 2**), in SCD mice it significantly increased mobilized CFU-C (2.6-fold) and showed a trend toward increased CFU-gmEM (2.1-fold, P=0.07) compared with vehicle-treated SCD mice (**Figures 4, B and C**). Importantly, four days of clotrimazole treatment further potentiated AMD3100-induced HSPC mobilization in SCD mice, leading to significantly higher CFU-C (1.9-fold) and CFU-gmEM (1.8-fold) compared with AMD3100 alone (**Figures 4, B and C**).

**Figure 4.**
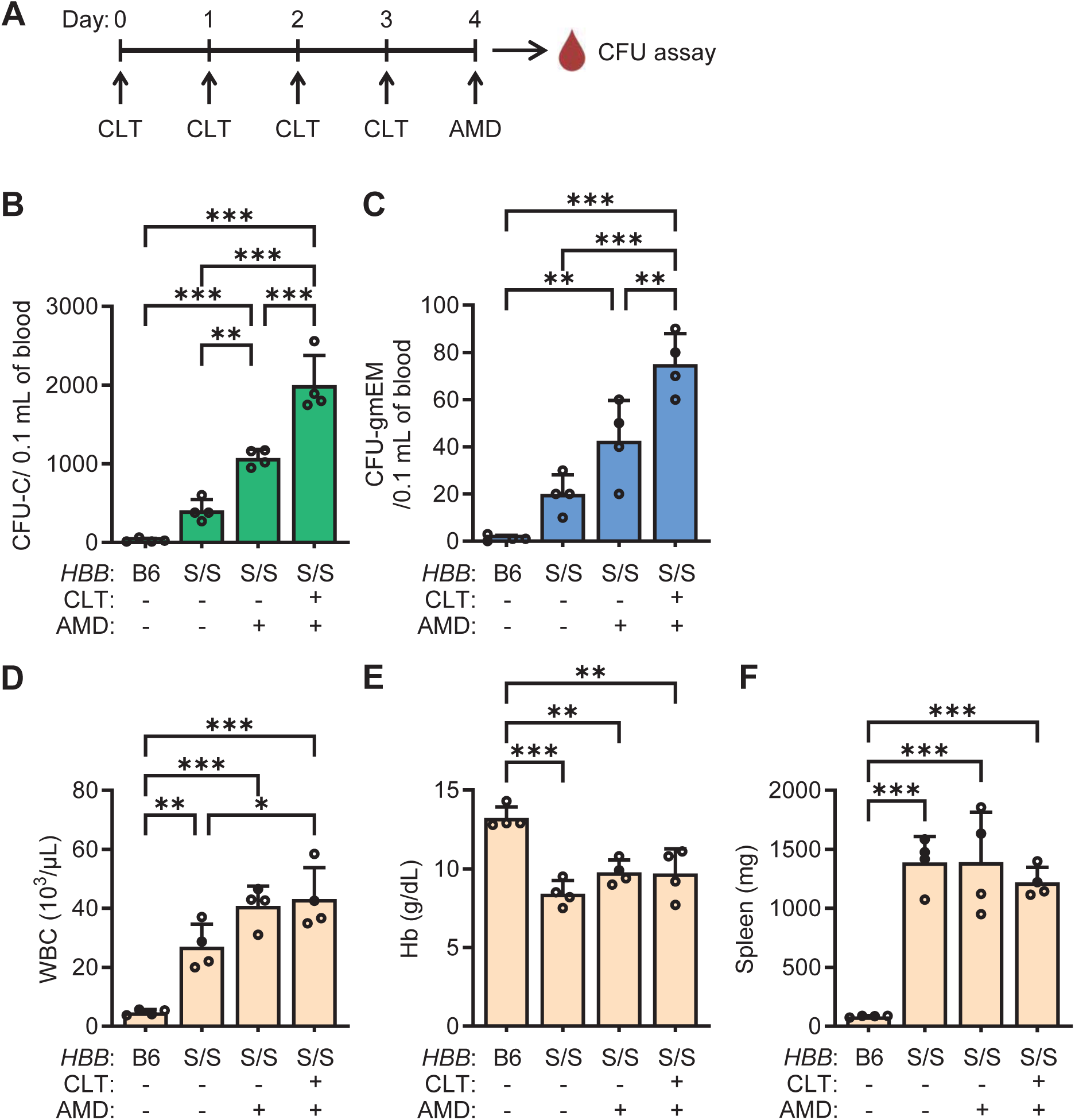
Clotrimazole enhances HSPC mobilization in sickle-cell disease mice in combination with AMD3100. (**A**) Experimental design for HSPC mobilization in “Townes” sickle-cell disease (SCD) mice. Mice were treated with CLT or vehicle daily for four days, followed by AMD3100 or vehicle one hour before peripheral blood collection for CFU assays. (**B**-**C**) Numbers of CFU-C (**B**) and CFU-gmEM (**C**) in peripheral blood after the indicated treatments (n=4 mice per condition from 4 experiments). B6, wild-type C57BL/6J mice; *HBB* S/S, mice homozygous for human sickle-cell β-globin alleles. (**D**-**F**) WBC counts, hemoglobin (Hb) levels, and splenic weights after the indicated treatments (n=4 mice per condition from 4 experiments). Data represent mean ± SD. Statistical significance was determined by one-way ANOVA with Tukey’s multiple comparisons test (^*^P<0.05, ^**^P<0.01, ^***^P<0.001).

As expected, SCD mice displayed elevated WBC counts relative to wild-type C57BL/6J mice (5.7-fold, **Figure 4D**). WBC counts were further increased by AMD3100 single treatment (1.5-fold, P=0.09) and by clotrimazole plus AMD3100 co-treatment (1.6-fold, P=0.04) compared with vehicle-treated SCD mice, but there was no significant difference between AMD3100 alone and co-treatment with clotrimazole (1.1-fold, P=0.97). SCD mice also exhibited reduced hemoglobin levels and splenomegaly compared with wild-type C57BL/6J mice, consistent with the disease phenotype; however, neither AMD3100 nor clotrimazole treatment altered these parameters (**Figures 4, E and F**). Taken together, these findings indicate that AMD3100 mobilizes HSPCs more effectively in SCD mice than in wild-type mice, and co-administration of clotrimazole further enhances this mobilization without exacerbating anemia or splenomegaly.

### Voriconazole enhances HSPC mobilization in wild-type and SCD mice

Although clotrimazole shows promising effects in enhancing HSPC mobilization, it is an FDA-approved only for topical and lozenge formulations and is not available for systemic use. To identify alternative Cyp7b1 inhibitors suitable for systemic administration, we tested voriconazole, another Cyp7b1-binding azole (29) that is FDA-approved for systemic antifungal therapy via both intravenous and oral routes. Importantly, voriconazole is already widely used in patients after HSPC transplantation as antifungal therapy (38), minimizing safety concerns regarding its potential repurposing for HSPC mobilization pre-transplant. In wild-type C57BL/6J mice, four days of intravenous voriconazole treatment significantly enhanced AMD3100-induced mobilization of CFU-C (1.5-fold) and CFU-gmEM (1.8-fold) into the peripheral blood (**Figures 5, A-C**). Similarly, in SCD mice, voriconazole co-treatment augmented AMD3100-induced mobilization, increasing CFU-C (1.5-fold) and CFU-gmEM (1.7-fold) compared with AMD3100 alone (**Figures 5, D and E**). These findings suggest that voriconazole, an FDA-approved systemic antifungal already in use among HSPC transplant recipients, could be rapidly repurposed as a clinically feasible strategy to enhance HSPC mobilization, including in patient populations such as SCD where current mobilization options are limited.

**Figure 5.**
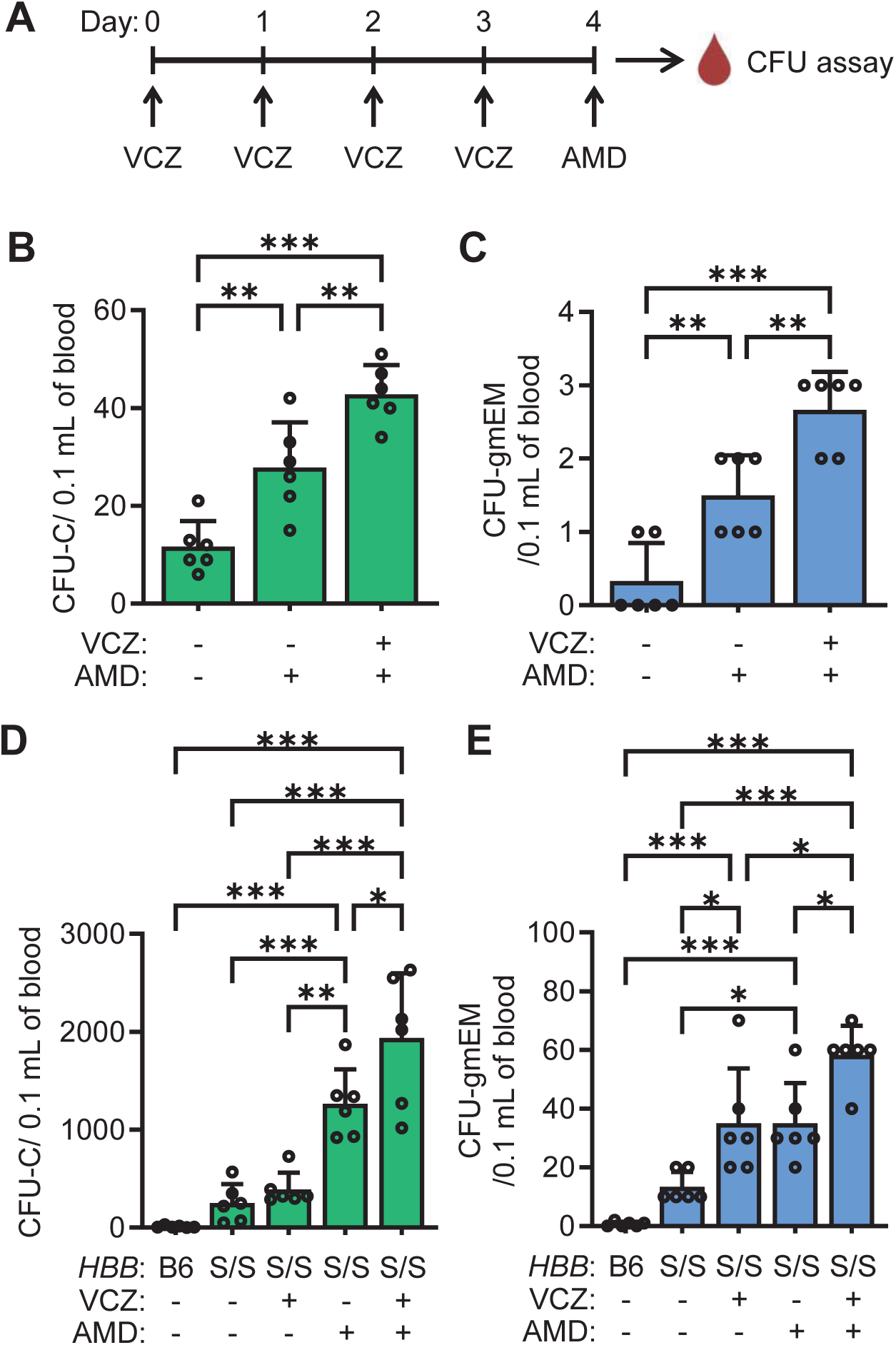
Voriconazole intravenous treatment enhances HSPC mobilization in wild-type and SCD mice in combination with AMD3100. (**A**) Experimental design for HSPC mobilization. Wild-type C57BL/6J or Townes SCD mice were treated with voriconazole (VCZ) or vehicle intravenously daily for four days, followed by AMD3100 or vehicle one hour before peripheral blood collection for CFU assays. (**B**-**C**) Numbers of CFU-C (**B**) and CFU-gmEM (**C**) in peripheral blood of C57BL/6J mice after the indicated treatments (n=6 mice per condition from 2 experiments). (**D**-**E**) The numbers of CFU-C (**D**) and CFU-gmEM (**E**) in peripheral blood of wild-type (B6) and SCD (*HBB* S/S) mice after the indicated treatments (n=6 mice per condition from 5 experiments). Data represent mean ± SD. Statistical significance was determined by one-way ANOVA with Tukey’s multiple comparisons test (^*^P<0.05, ^**^P<0.01, ^***^P<0.001).

### Voriconazole administration does not affect hematopoiesis

We next examined whether voriconazole administration alters hematopoiesis beyond its effect on mobilization. Following four days of intravenous voriconazole treatment in C57BL/6J mice, we quantified hematopoietic populations, including hematopoietic stem cells (HSCs), multipotent progenitors (MPPs), hematopoietic progenitor cells (HPC-1 and HPC-2), common myeloid progenitors (CMPs), granulocyte-macrophage progenitors (GMPs), megakaryocyte-erythroid progenitors (MEPs), granulocytes (Gra), monocytes/macrophages (Mac), erythroid progenitors (Ery), B cells, and T cells, in the bone marrow, spleen, and peripheral blood. Voriconazole treatment did not alter the cell numbers of any of these populations, nor did it affect WBC, red blood cell (RBC), hemoglobin, and platelet counts (**Figures 6, A-C**). Moreover, 5-bromo-2’-deoxyuridine (BrdU) incorporation assay during the last three days of treatment revealed no change in HSC cell-cycle activity in voriconazole-treated mice (**Figure 6D**).

**Figure 6.**
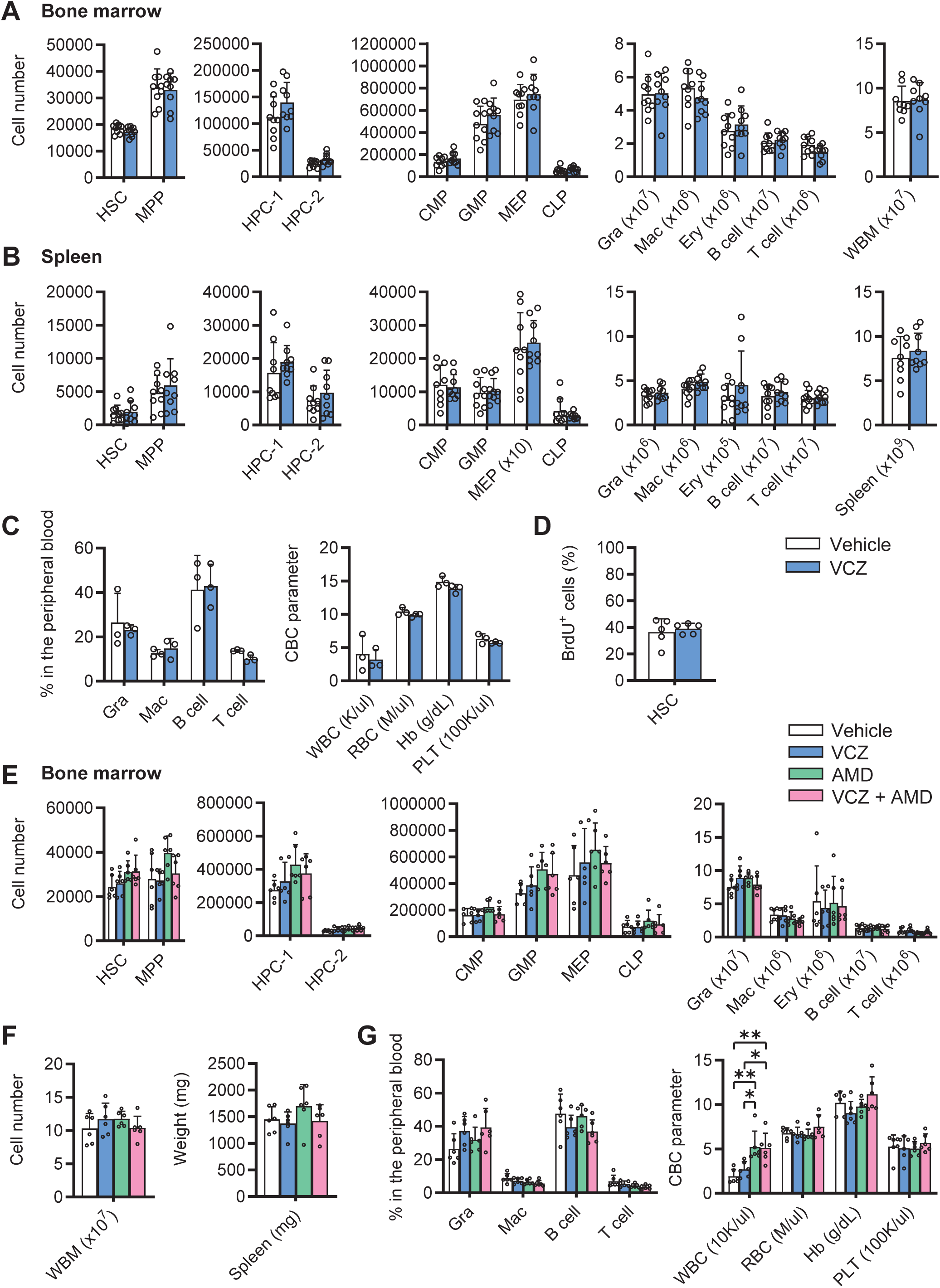
Voriconazole administration does not affect hematopoiesis. (**A**-**D**) Wild-type C57BL/6 mice treated with VCZ or vehicle intravenously daily for four days with BrdU incorporation during the last three days. (**A**-**B**) Flow cytometric quantification of hematopoietic populations in bone marrow (**A**) and spleen (**B**) (n=9 mice per condition from 3 experiments). (**C**) Frequencies of peripheral blood cell types (left) and complete blood count (CBC) parameters (right; Hb, hemoglobin; PLT, platelets) (n=3 mice per condition from 1 experiment). (**D**) Frequencies of BrdU^+^ cells within the HSC population (n=5 mice per condition from 2 experiments). (**E**-**G**) Townes SCD mice were treated with VCZ or vehicle intravenously daily for four days, followed by AMD3100 or vehicle one hour before analysis (n=6 mice per condition from 5 experiments). (**E**) Flow cytometric quantification of hematopoietic populations in bone marrow. (**F**) Bone marrow cellularity (left) and spleen weights (right). (**G**) Frequencies of peripheral blood cell type (left) and CBC parameters (right). Data represent mean ± SD. Statistical significance was determined by two-tailed unpaired Student’s t-tests (**A**-**D**) or one-way ANOVA with Tukey’s multiple comparisons test (**E**-**G**) (^*^P<0.05, ^**^P<0.01).

We also evaluated hematopoietic cell populations in SCD mice treated with vehicle, voriconazole alone, AMD3100 alone, or the combination of voriconazole and AMD3100. Aside from the expected increase in WBC counts observed in both AMD3100 alone and voriconazole plus AMD3100 treatment groups, no significant differences were detected in bone marrow hematopoietic populations (**Figure 6E**), whole bone marrow (WBM) cellularity and spleen weight (**Figure 6F**), or peripheral blood hematopoietic populations and complete blood count (CBC) parameters (**Figure 6G**) across treatment groups. Together, these findings indicate that the effects of voriconazole treatment are restricted to enhancing HSPC mobilization, without perturbing HSC cell-cycle dynamics, lineage differentiation, or peripheral blood composition. This selective activity underscores the safety of voriconazole treatment and supports its potential as a clinically viable mobilization agent in combination with AMD3100.

### Voriconazole augments but does not prolong HSPC mobilization

To determine whether the mobilization effect of voriconazole is transient or persistent, we examined the kinetics of mobilized HSPCs following co-treatment with voriconazole and AMD3100 in Townes SCD mice (**Figure 7A**).

**Figure 7.**
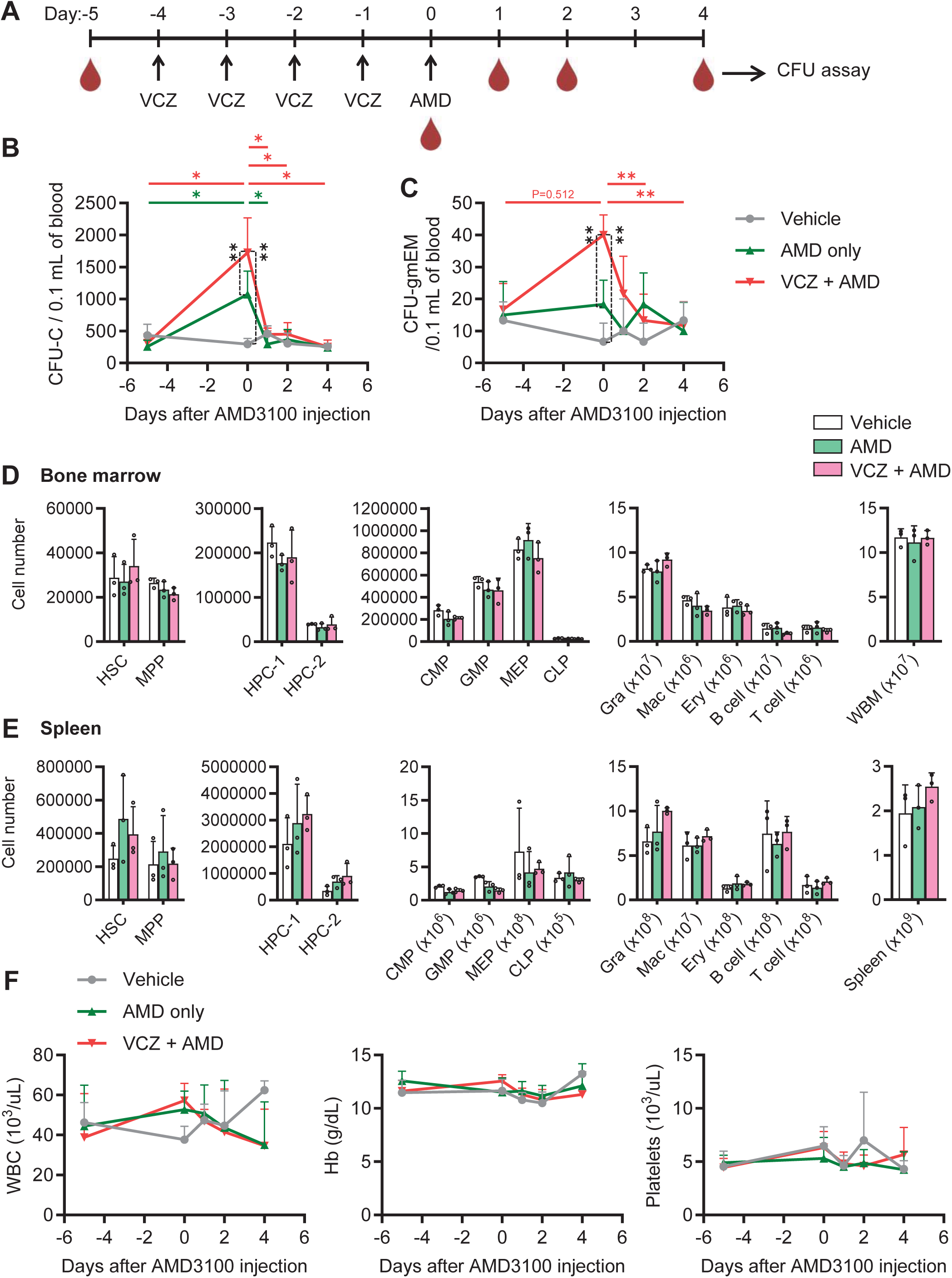
Voriconazole enhances HSPC mobilization transiently but not persistently. (**A**) Experimental design to study the kinetics of HSPC mobilization. Townes SCD mice were treated with VCZ or vehicle intravenously daily for four days, followed by AMD3100 or vehicle. Peripheral blood was collected for CFU assays and CBC on day-5, 1 hour, and days 1, 2, and 4 post-AMD3100 treatment. Bone marrow and spleen were harvested on day 4. (**B**-**C**) Numbers of CFU-C (**B**) and CFU-gmEM (**C**) in peripheral blood with the indicated treatments (vehicle: n=3 mice; AMD and VCZ+AMD: n=6 mice per condition from 2 experiments). (**D**-**E**) Flow cytometric quantification of hematopoietic populations in bone marrow (**D**) and spleen (**E**) (n=3 mice per condition from 1 experiment). (**F**) Kinetics of CBC parameters (vehicle: n=3 mice; AMD and VCZ+AMD: n=6 mice per condition from 2 experiments). Data represent mean ± SD. Statistical significance was determined by mixed-effects analysis with Šídák’s multiple comparisons test (**B**, **C**, and **F**) or one-way ANOVA with Tukey’s multiple comparisons test (**D** and **E**) (^*^P<0.05, ^**^P<0.01).

As expected, AMD3100 alone induced a significant increase in mobilized CFU-C one hour after injection, with levels returning to baseline within one day, consistent with prior reports (**Figure 7B**) (39). Co-treatment with voriconazole and AMD3100 further augmented CFU-C mobilization at one hour after AMD3100 injection compared with AMD3100 alone, but this effect was no longer evident after one day, when mobilized CFU-C numbers were comparable across vehicle-, AMD3100-, and combination-treated mice (**Figure 7B**). Mobilization kinetics of CFU-gmEM followed a similar pattern; however, a modest but nonsignificant increase persisted at one-day post-AMD3100 treatment, suggesting that voriconazole may exert a slightly more durable effect on immature subsets of HSPCs (**Figure 7C**).

Bone marrow hematopoietic populations remained comparable across treatment groups at four days post-treatment (**Figure 7D**). A trend toward increased splenic HSCs was observed in both AMD3100 alone (2-fold, P=0.32 by one-way ANOVA) and voriconazole plus AMD3100 (1.6-fold, P=0.61) cohorts, consistent with transient HSPC distribution to extramedullary sites (**Figure 7E**). No significant differences were observed in WBC counts, hemoglobin levels, or platelet counts between AMD3100 single and voriconazole co-treatment groups (**Figure 7F**). Together, these findings indicate that voriconazole enhances AMD3100-induced HSPC mobilization in a transient manner without perturbing steady-state hematopoiesis, further supporting its safety profile for potential clinical HSPC mobilization.

## Discussion

In this study, we identify a novel strategy for HSPC mobilization by targeting oxysterol metabolism using 27HC or Cyp7b1 inhibiting azoles (clotrimazole and voriconazole), in combination with clinically established HSPC mobilizers G-CSF and/or AMD3100. Notably, this strategy was effective not only in wild-type but also in SCD mice, where mobilization strategies are limited. This is clinically important because current FDA-approved gene therapies for SCD require substantially larger HSPC collections than standard autologous HSPC transplantation (7, 8), yet safe and efficient mobilization remains a major barrier. Importantly, we did not observe adverse effects of voriconazole on hematopoiesis in either wild-type or SCD mice. Voriconazole is already FDA-approved for systemic antifungal therapy and has been administered to patients undergoing HSPC transplantation (38). Our data reduce concerns about safety and raise the translational possibility that voriconazole could be repurposed to improve HSPC mobilization in clinical settings. In practice, patients may be able to receive additional short-course voriconazole dosing prior to HSPC collection to enhance mobilization efficiency.

Our findings suggest a model in which Cyp7b1 functions as a metabolic checkpoint in LT-HSCs. High expression of Cyp7b1 in LT-HSCs may restrict intracellular 27HC accumulation and thereby limit mobilization potential. Pharmacologic inhibition of Cyp7b1 could transiently release this brake, creating a mobilization-permissive state. While our previous study implicates ERα signaling as a key driver of this process (17), additional work will be needed to determine whether Cyp7b1 inhibition intersects with other pathways and whether it can act synergistically with newer mobilizers such as motixafortide (40), tGro-β (41), and VLA4 antagonists (42).

At the mechanistic level, Cyp7b1 catalyzes the hydroxylation of 27HC and 25HC into their downstream metabolites, 7α,27-dihydroxycholesterol (7α,27HC) and 7α,25-dihydroxycholesterol (7α,25HC), respectively (16). These metabolites have distinct signaling functions: 7α,27HC acts as an agonist of RAR-related orphan receptor gamma t (RORγt) (43), while 7α,25HC serves as a ligand for the GPR183 receptor (also known as EBI2) (44, 45). Although both 27HC and 25HC can signal through estrogen receptors and LXRs (15), our results show that only 27HC enhances HSPC mobilization, whereas 25HC does not. This indicates that HSPC mobilization is mediated specifically by 27HC signaling, likely through ERα as shown in our previous study (17). Accordingly, Cyp7b1-inhibiting azoles appear to enhance mobilization by blocking 27HC metabolism, rather than by modulating 25HC pathways. Moreover, because 27HC and related oxysterols may act on both HSPCs and their niche microenvironment, it will be important to determine whether enhanced mobilization reflects the combined contributions of cell-intrinsic and extrinsic mechanisms.

Beyond transplantation, enhanced mobilization by Cyp7b1 inhibitors may have broad applications, such as in vivo gene therapies. The current gene therapies FDA-approved for treating SCD involve ex vivo manipulation of HSPCs (7, 8). While this is a promising approach for treating several monogenic disorders, including immunodeficiencies, metabolic disorders, and osteopetrosis, it requires robust HSPC collection, maintaining HSPCs ex vivo during genetic modification, and patient conditioning for transplantation (46). Importantly, these procedures can promote positive selection of mutant clones harboring potential driver mutations associated with clonal hematopoiesis or myeloid neoplasms during ex vivo culture (47). In vivo gene therapy has the possibility to overcome these obstacles, and HSPC mobilization can enhance HSPC gene transfer in mouse models (48). Enhancing HSPC mobilization by Cyp7b1-inhibiting azoles therefore has the potential to directly improve the efficiency of in vivo HSPC gene therapy.

Mobilization strategies must also consider patient populations beyond SCD. For example, autologous HSPC transplantation remains a standard therapy for multiple myeloma (49), where chemotherapy-based HSPC mobilization has been used to provide both HSPC mobilization and anti-tumor activity. It will be important to determine whether elevated Cyp7b1 inhibition can further enhance chemotherapy-based HSPC mobilization while maintaining tolerability.

Finally, these results establish oxysterol metabolism as a novel regulator of HSPC mobilization. By targeting Cyp7b1 with clinically relevant agents such as voriconazole, it may be possible to develop mobilization strategies that are mechanistically distinct from, yet complementary to, current approaches. These strategies not only hold promises for expanding mobilization options in SCD patients, who cannot safely receive G-CSF, but also offer broader applicability across transplantation and gene therapy. Together, these findings position Cyp7b1 inhibition as a clinically translatable strategy with the potential to expand mobilization options and improve patient outcomes.

## Methods

### Sex as a biological variable

Mice of both sexes were used for the study, and similar results were observed.

### Animal studies

The C57BL/6J (RRID:IMSR_JAX:000664), B6;129-*Hbb^tm2(HBG1,HBB*)Tow^*/*Hbb^tm3(HBG1,HBB)Tow^ Hba^tm1(HBA)Tow^*/J (Townes model, RRID:IMSR_JAX:013071) (37), and NOD.Cg-*Kit^W-41J^ Tyr* ^+^ *Prkdc^scid^ Il2rg^tm1Wjl^*/ThomJ (NBSGW, RRID:IMSR_JAX:026622) (36) mice were obtained from The Jackson Laboratory. Young adult mice (2-3 months of age) were used. Mice were injected subcutaneously with 27HC (10 mg/kg/day, Avanti Polar Lipids) in 15% 2-hydroxypropyl-β-cyclodextrin (Sigma), G-CSF (250 μg/kg/day, filgrastim, Sigma) in phosphate buffered saline without calcium or magnesium (PBS, Corning), or AMD3100 (5 mg/kg, plerixafor, Sigma) in PBS, intraperitoneally with clotrimazole (50 mg/kg/day, Sigma) in corn oil (Sigma), or intravenously with voriconazole (50 mg/kg/day, MedChemExpress) in 30% 2-hydroxypropyl-β-cyclodextrin. All mice used in this study were housed in AAALAC-accredited, specific-pathogen-free animal facility at UConn Health.

### Cell and tissue preparation

Peripheral blood was collected from either submandibular vein, retro-orbital sinus, or caudal vena cava. Complete blood count was performed with the Vetscan HM5 Hematology Analyzer (Zoetis). Bone marrow cells were isolated by crushing the femurs and tibias with a mortar and pestle in staining medium (PBS without calcium and magnesium, supplemented with 1% heat-inactivated fetal bovine serum (Lonza)) and filtered through a 70-μm nylon screen (ELKO Filtering). Spleens were dissociated by crushing, followed by gentle trituration and filtering through a 70-μm cell strainer (Fisher Scientific). Bone marrow mononuclear cell number and viability were assessed by acridine orange/propidium iodide (AO/PI) staining (AO from Fisher Scientific; PI from Sigma) and counted by the LUNA-FL Dual Fluorescence Cell Counter (Logos Biosystems).

### Colony formation assay

Peripheral blood mononuclear cells from 10 uL to 100 uL of the blood, depending on the levels of HSPC mobilization, were isolated by Ficoll-Paque Premium density gradient media (1.085 g/mL density for mice and 1.078 g/mL density for humans, Cytiva) according to the manufacturer’s instructions. Isolated cells were seeded in a well of a 6-well plate containing Mouse or Human Methylcellulose Complete Media (R&D Systems) supplemented with 10 ng/ml mouse or human thrombopoietin (BioLegend). Colonies were counted after 12 days.

### Flow cytometry for mouse tissues

For isolation of HSCs and progenitors, a mixture of antibodies against CD2, CD3, CD5, CD8α, B220, Gr-1, and Ter119 was used to stain lineage markers as previously described (50). Red blood cells were lysed with ammonium-chloride-potassium lysing buffer. Non-viable cells were excluded during flow cytometry by staining with 4’,6-diamidino-2-phenylindole (DAPI, Tocris Bioscience). Antibodies (and clones) used in this study were anti-CD2 (RM2-5), anti-CD3 (17A2), anti-CD5 (53-7.3), anti-CD8α (53-6.7), anti-CD11b (M1/70), anti-CD16/32 (93), anti-CD31 (MEC13.3), anti-CD34 (RAM34), anti-CD45 (30-F11), anti-CD45R/B220 (RA3-6B2), anti-CD48 (HM48-1), anti-CD71 (R17217), anti-CD105 (MJ7/18), anti-CD117/c-kit (2B8), anti-CD135/Flt3 (A2F10), anti-CD150 (TC15-12F12.2), anti-Sca-1 (D7), anti-Gr-1 (RB6-8C5), and anti-Ter119 (TER-119). Antibodies and streptavidin were conjugated to one of the following dyes or to biotin: Brilliant Violet 421, Brilliant Violet 510, FITC, PerCP-Cy5.5, PE, PE-CF594, PE-Cy7, APC, Alexa Fluor 700, APC-Fire750, or APC-eFluor 780.

All antibodies were purchased from BioLegend, eBioscience, BD Biosciences, or Tonbo Biosciences. Data acquisition and cell sorting were performed using a FACSymphony S6 Cell Sorter, FACSymphony A5 SE Cell Analyzer, or LSR II Flow Cytometer (BD Biosciences) and data were analyzed using FlowJo v10 software (BD). The marker combinations used to identify mouse hematopoietic stem and progenitor cell populations examined in this study were:

HSC (hematopoietic stem cell): CD150^+^CD48^-/low^Lineage^-^Sca-1^+^c-kit^+^

MPP (multipotent progenitor): CD150^-^CD48^-/low^Lineage^-^Sca-1^+^c-kit^+^

HPC-1 (hematopoietic progenitor cell-1): CD150^-^CD48^+^Lineage^-^Sca-1^+^c-kit^+^

HPC-2 (hematopoietic progenitor cell-2): CD150^+^CD48^+^Lineage^-^Sca-1^+^c-kit^+^

CMP (common myeloid progenitor): CD34^+^CD16/32^low^Lineage^-^Sca-1^-^c-kit^+^

GMP (granulocyte-macrophage progenitor): CD34^+^CD16/32^high^Lineage^-^Sca-1^-^c-kit^+^

MEP (megakaryocyte-erythroid progenitor): CD34^-/low^CD16/32^low^Lineage^-^Sca-1^-^c-kit^+^

Gra (granulocyte): CD11b^+^Gr-1^+^B220^-^CD3^-^

Mac (monocyte/macrophage): CD11b^+^Gr-1^-^B220^-^CD3^-^

Ery (erythroid progenitor): CD71^+^Ter119^+^CD45^+^

B cell: B220^+^CD3^-^CD11b ^-^Gr-1^-^

T cell: CD3^+^B220^-^CD11b ^-^Gr-1^-^

### Cell-cycle analysis

To analyze BrdU incorporation rates in vivo, mice were given an intraperitoneal injection of 100 mg/kg of BrdU (Sigma) in PBS and maintained on 1 mg/ml BrdU in the drinking water for three days. Bone marrow cells were incubated with biotinylated antibodies against lineage markers, followed by MojoSort Streptavidin Nanobeads (BioLegend), and Lineage^+^ cells were depleted using a MojoSort Magnet (BioLegend). Frequencies of BrdU positive cells within the HSC compartment were measured by flow cytometry using the BrdU Flow Kit (BD Biosciences) with anti-BrdU antibody (3D4) conjugated with PE-Cy7 (BioLegend).

### Human cord blood HSPC xenotransplantation

Anonymous human cord blood samples were obtained from the Department of Obstetrics and Gynecology at UConn Health. CD34-positive cells from each donor were enriched by Ficoll-Paque Premium density gradient media (1.078 g/mL density) followed by EasySep Human CD34 Positive Selection Kit II (STEMCELL Technologies) according to the manufacturer’s instructions. Cell number and viability were assessed by AO/PI staining and counted by the LUNA-FL Dual Fluorescence Cell Counter. NBSGW recipient mice were transplanted with 5×10^4^ CD34-enriched cells via retro-orbital injection. Four months after transplantation, recipient mice were treated with mobilizing agents. Bone marrow cells were stained with anti-mouse CD45 (30-F11), anti-human CD45 (HI30), anti-HLA-ABC (W6/32), anti-human CD34 (581), and anti-human CD38 (HB7) antibodies. Peripheral blood cells are used for colony-forming assay. Derived colonies were stained with anti-mouse CD45 (30-F11), anti-human CD45 (HI30), anti-human CD33 (WM53), and anti-human CD235a (HIR2) antibodies.

### Statistics

Data are shown as mean ± standard deviation. The sample size used in each experiment was not formally justified for statistical power. No formal blinding was applied when performing the experiments or analyzing the data. Mice were allocated to experiments randomly and samples processed in an arbitrary order, but formal randomization techniques were not used. For analysis of the statistical significance of differences between two groups, we performed two-tailed unpaired Student’s t-tests. For analysis of the differences among more than two groups, we performed one-way ANOVAs with Tukey’s multiple comparisons test taking each cell population as one family. All statistical tests were performed using the GraphPad Prism software.

## Study approval

All animal procedures were approved by the UConn Health Institutional Animal Care and Use Committee.

## Data availability

All individual values represented in graphs are available upon request.

## Author contributions

HO conceived of the study and planned experiments. BLV and TJR performed most of the experiments with help from JKK, ALK, ET, and HO. BLV and HO analyzed data and wrote the manuscript. HO acquired funding and supervised the study.

## Acknowledgments

This work was funded by the National Blood Foundation and the National Institute of Diabetes and Digestive and Kidney Diseases (R01DK125747). We thank the UConn Health Flow Cytometry Facility. Illustrations were created with BioRender.

